# Stimulus sensitivity in noisy neural systems

**DOI:** 10.64898/2025.12.24.696343

**Authors:** Carlo Paris, Steeve Laquitaine, Matthew Chalk, Ulisse Ferrari

## Abstract

Understanding how neural populations encode sensory information requires a precise definition of neuronal sensitivity to stimuli. While tuning curves and firing rates offer intuitive insights, theoretical frameworks based on signal-to-noise ratio and decoding efficiency identify Fisher information as the canonical measure of sensitivity, due to its relation with decoding performance. However, this relation holds only under restrictive conditions—when many neurons encode the same stimulus feature or when neural noise is weak. In realistic settings, such as when complex or high-dimensional stimuli are represented by a small ensemble of neurons, Fisher information becomes ill-defined or misleading. To overcome these limitations, we investigate two complementary information-theoretic quantities—the stimulus-specific information (*I*_*SSI*_) and the local information (*I*_*loc*_)—and propose them as robust alternatives for quantifying sensitivity. We show that *I*_*SSI*_ and *I*_*loc*_ converge with Fisher information when signal to noise is large, yet remain meaningful and interpretable beyond that regime. Importantly, these measures capture distinct aspects of sensitivity: *I*_*SSI*_ quantifies how observing a response reduces stimulus uncertainty, whereas *I*_*loc*_ reflects how small stimulus perturbations reshape the system’s posterior beliefs. Together, they offer a unifying perspective linking information-theoretic and statistical notions of sensitivity, bridging theoretical analysis and experimental investigation of neural coding.

## 1 Introduction

Ever since Hubel and Wiesel first observed simple cells in V1 responding selectively to specific orientations and spatial locations (1; 2; 3), neuroscientists have sought to quantify which stimuli neurons are sensitive to. But what does it mean for a neuron to be sensitive to a stimulus? For a single neuron, this notion is relatively intuitive: we can measure how its firing rate changes as we vary a particular stimulus feature, such as the orientation or location of a bar of light, producing a tuning curve that captures its selectivity. The neuron is then said to be most sensitive to stimulus changes that elicit the largest change in firing rate—that is, where its tuning curve is steepest (4; 5; 6; 7). For example, a V1 simple cell responds strongly when a bar of light appears within a specific region of visual space, the receptive field, but weakly when the bar is moved elsewhere, indicating that it is sensitive to position within that region.

Extending this intuition from single neurons to even small populations of neurons, which jointly encode entire visual scenes, is less straightforward. Population activity lives in a multi-dimensional space, where the simple notion of a single “tuning curve” no longer applies, and it is unclear how to define where responses change most steeply. To make this notion precise, researchers often turn to the Fisher information (8; 9), which quantifies how strongly the responses of a neural population vary in response to small changes in the stimulus. The Fisher information provides a formal measure of how sensitive the neural code is to infinitesimal stimulus perturbations (10; 11; 12; 13). Further, the fact that it is often relatively straightforward to compute from neural responses has contributed to its widespread use in computational neuroscience (for a detailed review see (14)).

A major theoretical justification of the Fisher information as a measure of neural sensitivity comes from the fact that it provides a lower bound on how well a stimulus can be estimated from neural responses. However, as we show here, in many cases of interest—such as when neural responses are unreliable and/or when the stimulus has as more dimensions than the recorded neural population—the assumptions underlying this relation no longer hold. In these regimes, the link between the Fisher information and estimation error begins to break down, weakening its justification as a measure of neural sensitivity (15; 16; 14). Using data from retinal response to either moving bars (17; 18), or flashed natural images (19), we show that this breakdown can make Fisher information a misleading measure of neural sensitivity under naturalistic conditions.

To address this limitation, we explore two alternative measures of neural sensitivity: the stimulus-specific information (*I*_*SSI*_) (20; 5) and the local information (*I*_*local*_) (21). Both are grounded in information theory (22; 23) as stimulus-dependent decompositions of the mutual information, which quantifies the total information neurons encode about all stimuli. Crucially, these decompositions remain valid even in high-dimensional or noisy regimes, where Fisher information no longer tracks estimation accuracy. Yet the two measures capture distinct notions of sensitivity. The *I*_*SSI*_ quantifies how much observing a neural response to a given stimulus reduces uncertainty about that stimulus—linking sensitivity to what can be inferred from the response. *I*_*local*_, by contrast, quantifies how sensitive neural responses are to small changes in the stimulus, serving as a generalized, smoothed version of the Fisher information. Using examples from retinal coding, we show how these measures differ qualitatively in what they reveal about neural sensitivity, and discuss when each is most appropriate depending on the aspect of information processing one wishes to capture.

Together, these results build on classical approaches to measuring neural sensitivity, providing tools that remain valid under realistic, high-dimensional stimulus conditions. They offer a practical framework for understanding how sensory systems encode the rich and variable structure of the natural world.

## 2 Results

### Fisher Information and the Cramér–Rao Bound

Firing rates, tuning curves, and receptive fields provide an intuitive notion of stimulus sensitivity: neurons appear most sensitive to stimuli that elicit the largest responses. This intuition, however, has drawbacks. A neuron that responds equally strongly to all stimuli would be assigned high sensitivity everywhere, despite conveying no information about which stimulus was presented. Moreover, response magnitude alone ignores neural variability: a stimulus that evokes a large but highly unreliable response can be far less informative than one producing a smaller yet consistent response. Sensitivity, should therefore depend not on the mean activity alone, but on how reliably responses distinguish nearby stimuli.

Alternatively, we might say that a neuron is “sensitive” to a stimulus, *x*, when small changes in the stimulus produce large and reliable changes in its response, *r*. Because neural responses are inherently noisy—the same stimulus rarely evokes identical responses—we can formalize this intuition by describing responses probabilistically, through the conditional distribution *p*(*r* | *x*). The *Fisher information* quantifies how strongly, on average, small variations in *x* alter this distribution:

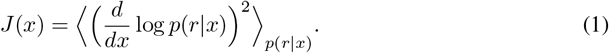

Large Fisher information means that infinitesimal stimulus changes produce large, systematic shifts in the response distribution.

Fisher information also connects naturally to classical *tuning curves, f* (*x*), which describe neuronal mean firing rates. For a single neuron encoding a one-dimensional feature, it obeys the inequality

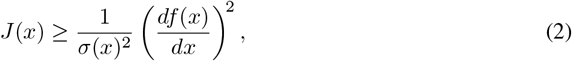

where *σ*(*x*)^2^ = Var(*r* | *x*) is the response variance. Thus, the Fisher information increases when the tuning curve changes steeply with *x* relative to the trial-to-trial noise. This also extends to multiple neurons: when neurons encode *x* independently, their Fisher informations simply add.

A fundamental theorem, the *Cramér–Rao bound* (CR bound) (24; 25), gives Fisher information a concrete operational meaning as a measure of sensitivity. The CR bound states that the mean-squared error of any unbiased estimate of the stimulus is bounded from below by the inverse of the Fisher information:

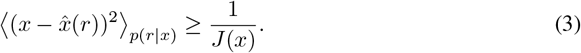

where 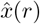 is an unbiased decoder of the neural activity, defined such that 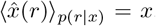. No unbiased decoder can therefore estimate *x* with a precision exceeding this limit. In other words, the Fisher information sets a theoretical bound on how accurately, in principle, small stimulus changes can be discriminated from noisy neural responses—the larger the Fisher, the finer the achievable discrimination (26; 23; 12).

However, the CR bound defines a lower bound on the decoding error only for unbiased decoders. In the case of vanishing or small Fisher information such a decoder can not be realized in practice (16), and the lower bound can be violated as we will show later. Generalisations of the Cramér–Rao bound, which apply to biased decoders, introduce additional quantities—notably the bias and its first derivative—so that decoding performance is no longer determined by the Fisher information alone. As a result, Fisher information by itself does not predict the performance of any realizable decoder, weakening its justification as a practical measure of stimulus sensitivity.

To test when the Fisher predicts the decoding performance, we simulated populations of neurons encoding motion direction with bell-shaped tuning curves (Fig.1A; see Methods). For a large population of neurons, the Fisher information closely matched the precision 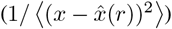 of an optimal Bayesian decoder, indicating that the CR bound was tight under these conditions. Reducing the population to only four neurons revealed a different picture, however. At both high and low signal-to-noise ratios (SNR), the Fisher information and decoding precision varied strongly across orientations (Fig. 1B–C), as individual neuronal contributions no longer averaged out to produce a smooth curve. Crucially, the close match between the Fisher information and the precision, that we observed in the larger population, completely disappeared in these conditions. At low SNR, we not only observed violation of the CR bound (Eq. 3), but the two measures were even anticorrelated: Fisher information peaked at the tuning-curve flanks, while decoder precision peaked near the preferred orientations (Fig. 1C, middle). For a single neuron (Fig. 1D), this discrepancy became extreme – Fisher information was large at the flanks, yet decoding precision increased only near the tuning-curve peak. Thus, the link between Fisher information and decoding precision can completely break down in small or noisy populations.

**Figure 1.**
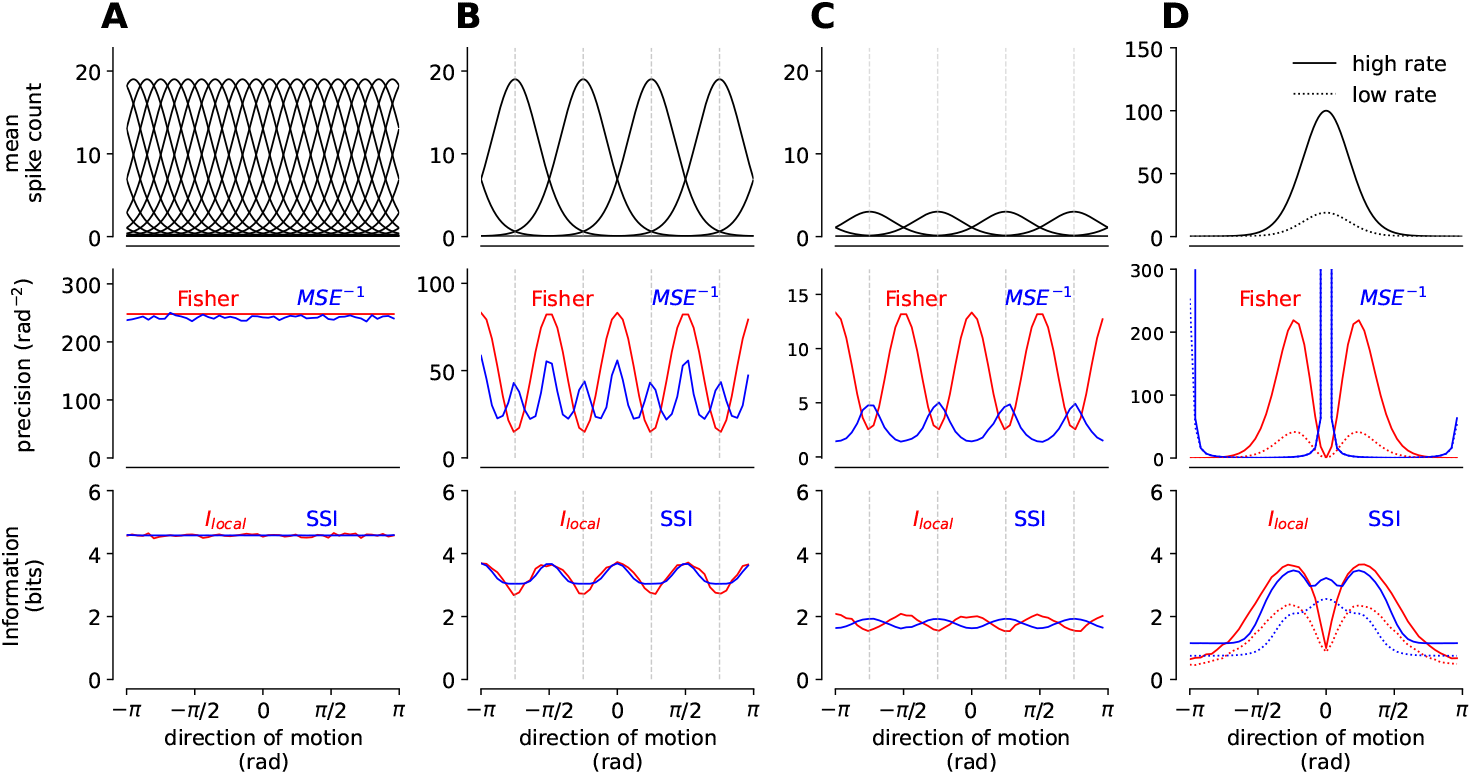
Fisher information can strongly differ from Bayesian decoder performance. **A.** Simulated responses of 20 model neurons of the Middle Temporal Area (MT) with homogeneous direction tuning to a moving stimulus. The top panels show representative Von-Mises shaped tuning curves. Motion directions are uniformly distributed. The middle panels compare the Fisher information (red) with the decoding precision (inverse mean-squared error, blue) from an optimal Bayesian decoder, demonstrating close agreement when the population is large and signal-to-noise ratio (SNR) is high. **B-C**. For smaller populations (4 neurons) and B) high-versus C) low-SNR conditions, Fisher information and precision diverge markedly, indicating that Fisher information no longer tracks decoder performance. **D**. Single-neuron case showing that Fisher information peaks at the tuning-curve flanks whereas decoding precision peaks at the tuning maximum, further illustrating the breakdown of the Fisher-Cramér-Rao link at low SNR. Bottom panels show stimulus-specific information (*I*_*SSI*_, blue) and local information (*I*_local_(*x*), red), which remain well-behaved under these conditions.

Next, we asked whether this scenario, where the Fisher information is uncoupled from decoding precision, occurs in realistic settings, where neural tuning curves are heterogeneous and irregularly cover the stimulus space. To test this we considered small populations of four On-Off direction selective retinal ganglion cells such that, within the quadruplet, each cell preferentially responded to a different cardinal direction (27; 28). The dataset consisted of electrophysiological recordings previously collected from the rabbit retina (17; 18; 29). We fitted their tuning curves with Flat-Topped Von Mises functions, an extension of the gaussian modified for rotational data (30). Fig. 2 shows four example quadruplets taken from this dataset, where 4 neurons jointly encode motion direction in a particular spatial location (see Methods). In Fig. 2A-B we see that, despite the fact there are only 4 neurons, when their firing rates are high enough, there is a tight relation between the Fisher information and the precision of a Bayesian decoder. In contrast, for quadruplets with lower firing rates, this coupling breaks down, with the Fisher information even anti-correlating with the precision of the Bayesian decoder in certain cases (e.g. Fig. 2D).

**Figure 2.**
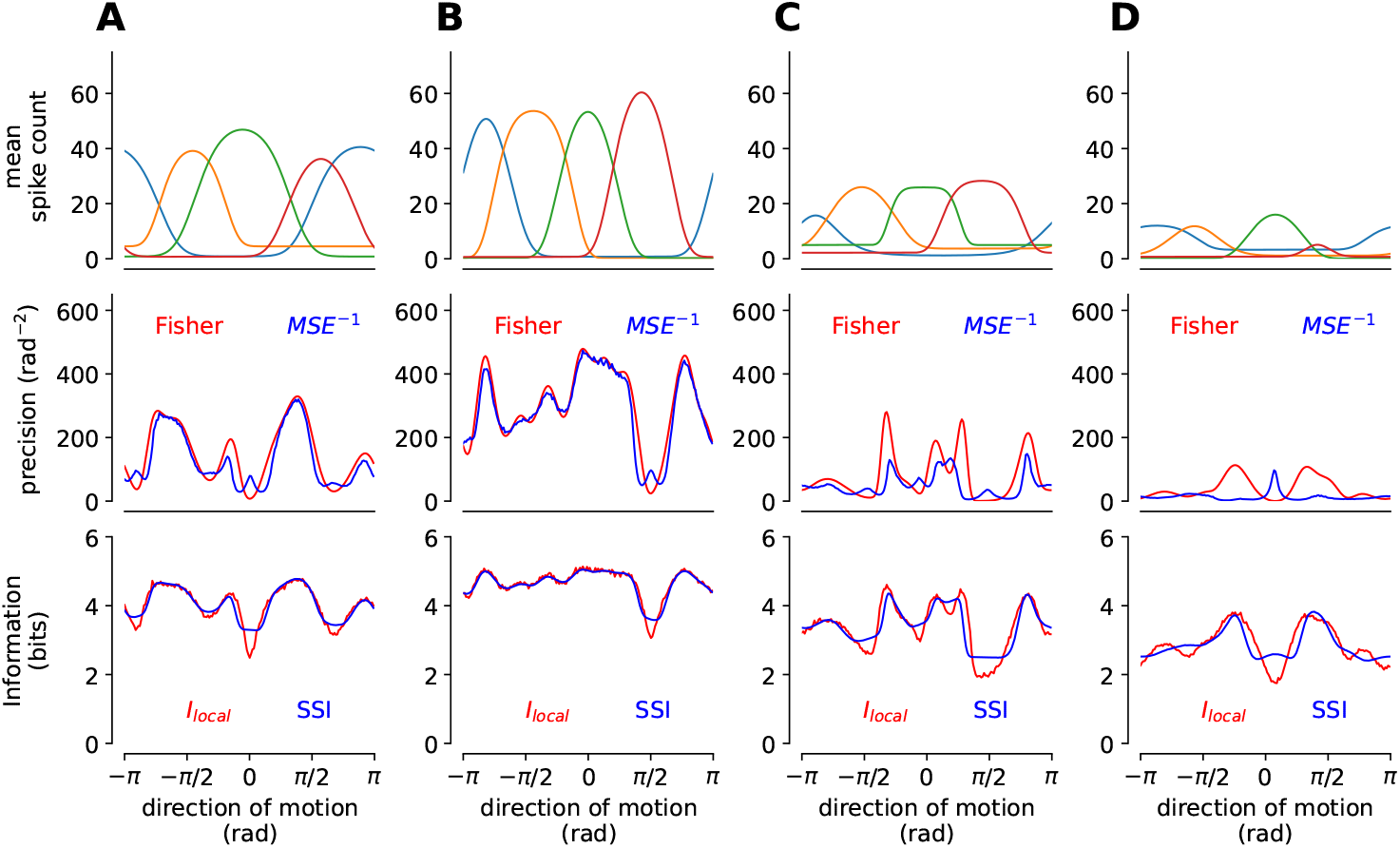
Sensitivity and decoding error for quadruplets of direction sensitive retinal ganglion cells (DS RGCs). **A-D.** Four examples of quadruplet tuning curves (top row). Comparison of Fisher information and inverse mean squared error (middle row), and of stimulus-specific information (*I*_*SSI*_) and local information (*I*_*local*_) (bottom row), across motion direction. These measures reveal distinct patterns of neural sensitivity: while Fisher and MSE-based precision can diverge under realistic noise conditions, both *I*_*SSI*_ and *I*_*local*_ remain consistent and interpretable, highlighting their robustness to variability in firing rate and tuning width.

How to understand this mismatch between decoding precision and Fisher information? As explained before, one reason is that the Cramér-Rao bound only holds strictly for *unbiased* decoders, where 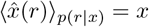. In many realistic neural settings, however, unbiased decoding is either unattainable or highly suboptimal. In our examples, both Bayesian and maximum-likelihood decoders became biased when the number of neurons was small or the SNR was low. In fact, when the Fisher information vanishes (e.g., for a single neuron whenever *f* ^*′*^(*x*) = 0; Fig. 1D), no locally unbiased estimator exists (16). Under such conditions, the Cramér-Rao bound no longer characterizes achievable performance, severing the link between Fisher information and optimal decoding error.

A second limitation arises because Fisher information is inherently *local*: it only depends on how responses change for infinitesimal perturbations of the stimulus around a point *x*. In contrast, optimal decoding - especially at low SNR or with small neural populations - depends on the *global* geometry of the likelihood *p*(*r* | *x*) and the prior *p*(*x*): the shape of response distributions at distant stimuli affects how confusable different stimuli are. When decoding relies on these global features, the Cramér-Rao bound can become loose even in the absence of bias, and Fisher information may no longer track true discriminability or estimation precision.

In short, Fisher information only captures local response sensitivity and assumes ideal, unbiased decoders—assumptions that often break down in realistic neural settings, especially with small populations or low SNR.

### Beyond Fisher: alternative measures of neural sensitivity

We next sought alternative measures of sensitivity that remain meaningful in cases with small neural populations and/or low noise, where the Cramér-Rao bound is loose or violated.

#### Local information

Fisher information measures how responses change for tiny stimulus shifts. This is powerful when responses are precise, but when noise is high - for example with small neural populations or low SNR - optimal decoding depends on the global structure of the response distribution, and Fisher no longer predicts actual decoding accuracy.

To move beyond infinitesimal perturbations and capture sensitivity over a finite range of stimulus variations, we introduce the *local information* (21). Rather than probing responses to an infinitely small change in the stimulus, we instead consider finite changes by computing how much information the observed responses carry about a *noisy* version of the stimulus,

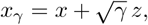

where *z* ∼ 𝒩 (0, 1) and *γ* controls the noise amplitude. We then evaluate the Fisher information with respect to this noisy stimulus, *J* (*x*_*γ*_), and integrate over all noise levels:

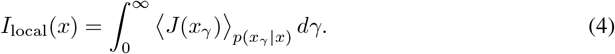

This procedure effectively smooths the Fisher information over multiple scales, yielding a measure of sensitivity that reflects how neural responses encode perturbations of all magnitudes, not just infinitesimal ones.

The local information is directly linked to Shannon’s mutual information *I*(*R*; *X*), which measures the total information neural responses convey about the full stimulus ensemble. In fact, the mutual information is simply the average of the local information over stimuli:

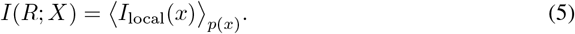

As we showed previously (21), *I*_local_(*x*) inherits key interpretability properties of the Fisher information: it is always non-negative, and - crucially - it remains *local*, in the sense that it depends primarily on the responses to stimuli in a neighborhood around *x*.

#### Stimulus-specific information

An alternative way to quantify sensitivity is to measure how much observing a neural response *r* reduces uncertainty about the stimulus *x*. Entropy *H*(*X*) captures this uncertainty: high when many stimuli are plausible, lower when the stimulus distribution is concentrated.

Following Butts (20; 5), we define the *stimulus-specific information* (*I*_*SSI*_) as the expected reduction in stimulus entropy after observing the response evoked by stimulus *x*:

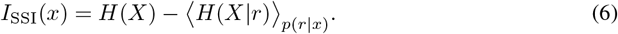

Stimuli whose responses strongly constrain the posterior *p*(*x* | *r*) have high *I*_*SSI*_; those that produce ambiguous responses have low *I*_*SSI*_. As with the local information, averaging *I*_SSI_(*x*) over *p*(*x*) yields the mutual information:

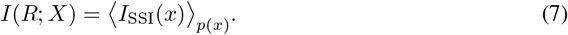

In contrast to *I*_local_(*x*), which characterises the sensitivity of neural responses to stimulus perturbations, *I*_SSI_(*x*) quantifies how much the neural responses evoked by stimulus *x* reduce a downstream observer’s uncertainty about which stimulus was presented (20; 11; 29). As such, *I*_SSI_(*x*) is naturally aligned with downstream decoding or inference tasks, where the goal is to identify the stimulus from sensory responses, rather than to assess local discriminability in stimulus space. On the other hand, unlike *I*_*local*_(*x*), *I*_*SSI*_ (*x*) is not strictly ‘local’, and thus it may be influenced by how neurons respond to distant stimuli, far from *x*.

#### Comparing alternative measures of sensitivity

To illustrate how these measures behave, we examined orientation coding in our simulated neural population. With many neurons, both *I*_local_ and *I*_SSI_ were uniform across orientations and equal to the mutual information *I*(*R*; *X*) (Fig. 1A, bottom panel). As with Fisher information, reducing the population size introduced stimulus-dependent structure (Fig. 1B–C). At high SNR, *I*_local_ and *I*_SSI_ aligned, peaking at the tuning-curve flanks (Fig. 1B), consistent with the intuition that these regions support fine discrimination. At low SNR, however, the measures diverged: *I*_local_ continued to peak at the flanks, while *I*_SSI_ shifted to peak at the tuning-curve centre (Fig. 1C). The same pattern appeared for a single neuron (Fig. 1D). This transition in *I*_SSI_ was described by Butts and Goldman (5): when noise is high, fine discrimination is not possible and responses near the tuning peak carry the most information. In contrast, *I*_local_ behaves as a smoothed Fisher information and consistently emphasizes tuning-curve flanks across SNR regimes. In short, *I*_SSI_ reflects how much uncertainty the responses actually reduce—strongly shaped by noise level and population size—whereas *I*_local_ tracks local sensitivity and therefore retains Fisher-like behaviour even when decoding performance shifts.

We observed a similar pattern when applying these sensitivity measures to recordings from quadruplets of direction-selective retinal ganglion cells (Fig. 2). When firing rates were high and the Fisher information closely tracked decoding precision (Fig. 2A–B), *I*_local_ and *I*_SSI_ were strongly correlated with each other and with the Fisher information. At lower firing rates (Fig 2C-D), however, this correspondence weakened, and we again found cases in which *I*_SSI_ peaked at the tuning-curve centre while *I*_local_ consistently peaked at the flanks (e.g. Fig. 2D). Thus, the intuition developed in our simulations carries over to neural data with heterogeneous, irregular tuning curves.

### Quantifying sensitivity with naturalistic stimuli

Low-dimensional toy stimuli like bars and gratings are useful, but real neurons encode high-dimensional natural scenes. To apply our approach to realistic settings, we extend our analysis to high-dimensional inputs.

For multidimensional stimuli, the Fisher information becomes a matrix:

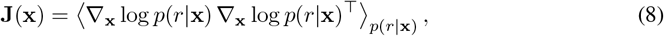

where ∇_**x**_ denotes the gradient with respect to the stimulus components. Likewise, the mean-squared decoding error is also a matrix, and the multivariate Cramér-Rao bound states:

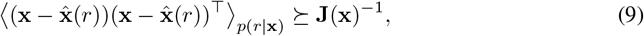

meaning that, for any direction **u** in stimulus space,

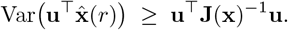

However, in many realistic settings the stimulus dimensionality exceeds the number of recorded neurons. This is common biologically (only a few ganglion cells tile each point in the retina) and experimentally (recordings capture only a tiny subset of neurons). In such cases, the Fisher matrix is necessarily low-rank and its inverse diverges, making the Cramér-Rao bound uninformative: no locally unbiased decoder exists, exactly as in the 1D case when the Fisher information dropped to zero (Fig. 1D). In simple cases, ignoring certain stimulus dimension might solve the issue (31), but for highly non-linear neurons the problem persists. Thus, classical Fisher-based bounds break down in high dimensions, motivating sensitivity measures that remain meaningful under naturalistic, high-dimensional stimulation.

In contrast, both *I*_SSI_ and *I*_local_ are grounded in Shannon mutual information, which remains well defined even when the stimulus space is high dimensional, discrete, or larger than the number of recorded neurons. These measures therefore continue to quantify sensitivity in regimes where the Cramér–Rao bound breaks down.

To examine their behaviour under naturalistic stimulation, we applied our framework to retinal ganglion cell responses evoked by natural images. Although the raw visual input was high dimensional, the neural responses were well described by a projection onto a two-dimensional stimulus subspace (see Methods). Importantly, even after this reduction, the stimulus dimensionality still exceeded the number of neurons analysed simultaneously, meaning that the Fisher information for individual cells is low-rank and the Cramér-Rao bound does not provide a meaningful constraint on decoding performance in this space.

These stimulus features, along with their natural-image prior distribution, are shown in Fig. 3 (top row). Across the dataset, tuning functions varied widely: some cells showed approximately monotonic, “simple-cell–like” responses, whereas others displayed more complex, quadratic-like tuning. For simple cells with high maximum firing rates (Fig. 3A), both *I*_local_ and *I*_SSI_ increased monotonically with the response, consistent with high-SNR behaviour. When firing rates were low (Fig. 3B), however, *I*_SSI_ changed shape and became large for stimuli that evoked little or no spiking—reflecting the fact that silence can be informative—while *I*_local_ remained largely unchanged. For cells with complex, non-monotonic tuning (Fig. 3C), the two measures diverged even more strongly: *I*_SSI_ peaked for stimuli producing little spiking, whereas *I*_local_ increased with firing rate.

**Figure 3.**
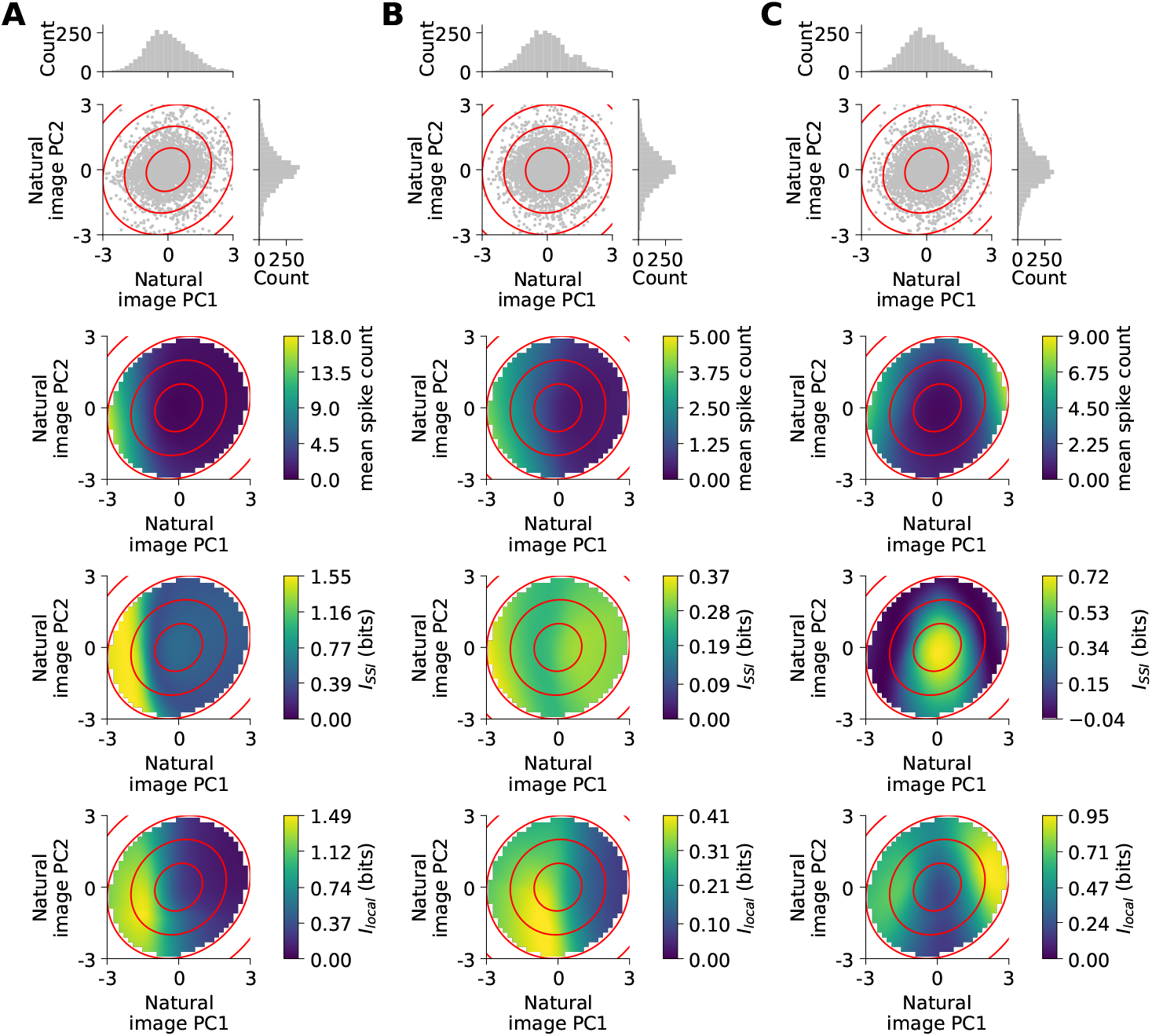
Sensitivity of retinal ganglion cells encoding natural images. (**A–C**) Distribution of natural images projected onto their first two principal components (PC1, PC2; top row). Mean firing rate of the neuron across the projected stimuli (second row). Stimulus-specific information (*I*_*SSI*_) and local information (*I*_*local*_) maps (third and fourth rows, respectively). Both *I*_*SSI*_ and *I*_*local*_ reveal structured sensitivity patterns aligned with the neuron’s tuning curve, illustrating their capacity to capture tuning-consistent variations in sensitivity even when Fisher information fails.

In summary, natural-image responses reproduce the pattern seen in our previous 1D simulations: *I*_SSI_ can change qualitatively with noise and tuning structure, while *I*_local_ behaves more consistently, acting as a smoothed Fisher information that emphasizes local sensitivity.

## 3 Discussion

This paper’s primary focus is the evaluation of neural sensitivity to stimuli. We explored three specific cases: first, a toy model with homogeneous tuning curves; second, tuning curves from On-Off DS RGCs exhibiting regions of low curvature and sparsely covering the stimulus ensemble; third, RGCs responses to natural images for which the effective dimensionality of the stimulus exceeds the number of neurons encoding it. We found that in all three cases the Fisher information fails to provide a lower bound for the decoding error and often violates the Cramér-Rao bound. Furthermore, we showed that the *I*_*SSI*_ and the *I*_*local*_ provide interpretable substitutes whose aptness depends on the chosen aspect of sensitivity one chooses to study.

The Fisher information has been often used in neuroscience, from psychophysics and detection theory (32; 10; 33), to the study of neural systems’ tuning properties (34), to population encoding (35; 36; 11) to its connection with information theory (37; 38), information limiting correlations (39; 40) and information geometry (13; 41; 12). However, a number of these studies based their metrics and conclusions on the Cramér-Rao bound, which relates the Fisher information to the mean squared decoding error. However, our results show how this simple relation does not hold in many biologically relevant cases and therefore cannot be uncritically relied upon. This does not necessarily invalidate previous findings, but highlights the importance of verifying the applicability of the bound when interpreting or extending such analyses.

Previous work has shown that the Cramér–Rao bound can be used to derive a close relationship between Fisher information and mutual information (37; 38). This relationship has subsequently been exploited to make predictions about how neurons encode stimuli so as to maximise mutual information under resource constraints (42; 43; 44; 45; 46; 47). These predictions were further extended to psychophysical performance, yielding quantitative biases and scaling laws that were later validated experimentally. However, because these results rely on the tightness of the Cramér–Rao bound, they are expected to break down in regimes where the (unbiased) bound does not hold, such as at high noise. It would therefore be interesting to ask when and how the predicted psychophysical biases begin to deviate from these theories as the Cramér–Rao bound ceases to be tight, for example when the visual signal is weak and neural responses become unreliable.

The choice of *I*_*SSI*_ versus *I*_*local*_ depends on the specific notion of sensitivity that one chooses to investigate, given their different interpretations. The primary strength of *I*_*SSI*_ lies in its interpretability: it’s a measure that quantifies how unambiguous a stimulus is for a downstream observer. Moreover, *I*_*SSI*_ ‘s behavior provides valuable insights into the sensory system’s encoding regime (5). This is particularly evident in the bottom panels of Fig. 1B-C: the *I*_*SSI*_ peaks at the tuning-curve flanks for high firing rates (B), corresponding to low-noise or slope encoding; and at the tuning curve peaks for low firing rates (C), corresponding to high-noise or peak encoding. However, the *I*_*SSI*_ is inherently non-local: its value at a given stimulus depends on the system’s responses to all other stimuli. This interdependence complicates intuitive predictions of its profile from the shape of the tuning curve alone. By contrast, the *I*_*local*_, though less directly interpretable, often mirrors the behavior of the Fisher information and is therefore more amenable to *a priori* evaluation. As illustrated again in Fig. 1B–C, the *I*_*local*_ closely tracks the Fisher information, with its minima and maxima aligning with those of the first derivative of the proximal tuning curve.

Our sensitivity measures, *I*_*SSI*_ or *I*_*local*_, also have some limitations: first, computing the *I*_*SSI*_ or *I*_*local*_ is considerably more demanding than calculating the Fisher information. In practice, these computational challenges can be addressed through the use of Monte Carlo methods for estimating the *I*_*SSI*_ (11; 29) and diffusion-based models for evaluating the *I*_*local*_ (21). Even if the link between Fisher information and decoding performance is weaker than one might expect, we have not found any such connection for neither *I*_*SSI*_ nor *I*_*local*_. For completeness, we note that the first can be connected with a lower bound on the posterior variance of a Bayesian decoder, which, however, might be very different from the actual decoding error.

Several studies have demonstrated that Fisher information can be misleading for small neural populations or limited data samples (11; 48; 49). Yarrow et al. (2012) (11) rigorously compared Fisher information and stimulus-specific information (*I*_*SSI*_) as metrics for assessing which stimuli are best encoded by a finite neural population: the peak of the tuning curve (indicative of high noise) or the flanks (low noise). They found convergence between the two metrics in large populations, where high SNR ensures robust, low-noise encoding. However, whereas the Fisher information consistently predicted maximal discriminability at the tuning curve flanks, the actual profile of single-neuron *I*_*SSI*_ shifted with population size - favoring peak or flank encoding depending on circuit size and noise - highlighting nuanced differences that only vanish for large *N*. These findings align with our results (Fig. 1), but Yarrow et al. focused on populations with homogeneous tuning curves and one-dimensional stimulus spaces, leaving the effects of heterogeneous tuning (Fig. 2) or multi-dimensional encoding (Fig. 3) largely unexplored. Taken together, these considerations underscore that while Fisher information offers a powerful summary for large and homogeneous populations, information-theoretic measures such as *I*_*SSI*_ and *I*_*local*_ may provide a more nuanced and robust framework for quantifying stimulus encoding, particularly in the heterogeneous or data-limited regimes characteristic of biological populations.

Finally, the link between Fisher information and sensitivity via decoding error breaks down when the stimulus dimensionality exceeds the number of encoding neurons. In contrast, both *I*_*SSI*_ and *I*_*local*_ provide well-defined measures whose relation to the mutual information remains valid regardless of the dimensionality of either stimulus or neural response spaces. This is important because, although artificial stimuli have been widely used due to their simple statistics and amenability to experimental control, a growing body of work suggests that stimuli statistics heavily impact neuronal responses (50; 51; 52). Thus, probing sensory circuits with naturalistic stimuli is likely to reveal fundamentally different functional characterizations compared to those obtained with conventional artificial inputs.

## 4 Methods

### Simulated populations of motion direction–selective neurons

We simulated motion direction tuning curves, *f* (*x*), representing the mean spike counts for each stimulus direction *x*, in neural populations comprising twenty, four, or a single neuron (Fig. 1). Each tuning curve was modeled using a von Mises function, parameterized after tuning properties observed in the middle temporal (MT) area of mammals (53; 54). Neurons had a maximum mean firing rate of 19 spikes and a concentration parameter *k* = 0.29, corresponding to a full width at half maximum (FWHM) of 75^°^ (53). Stimulus direction *x* was expressed in radians within the interval [0, 2*π*], and preferred directions were uniformly distributed across the stimulus space. Low-, intermediate-, and high-SNR conditions were tested using maximum mean spike counts of 3, 20, and 100 spikes, respectively.

The prior distribution of motion directions, *p*(*x*), was assumed to be uniform, while the response likelihoods were assumed to be conditionally independent Poisson distributions.

### Model of direction selectivity in Retinal Ganglion Cells

We selected four representative examples of quadruplets of retinal ganglion cell (RGC) tuning curves, *f* (*x*), obtained by fitting the RGCs’ mean responses to motion in different directions (18; 17), using Flat-Topped Von Mises functions (30). Each quadruplet was heterogeneous, with tuning curves differing in width, preferred motion direction, maximum mean spike count, and baseline firing rate. As in the simulated motion direction–selective populations, motion direction *x* was expressed in radians over the interval [0, 2*π*].

The prior distribution of motion directions, *p*(*x*), was assumed to be uniform, while response likelihoods were assumed to be conditionally independent Poisson distributions.

### Model of Retinal Ganglion Cell responses to natural images

Responses of 41 a mouse retinal ganglion cells (RGCs) to 3,160 natural images were analyzed (55; 19). Principal component analysis (PCA) was applied to the images, and the first two principal components were retained. Each neuron’s two-dimensional tuning function, *f* (**x**), where **x** denotes the vector of PC_1_ and PC_2_ values for each image, was modeled using a deep neural network trained to predict RGC responses from these components.

The fully connected feedforward neural network had the following architecture:

- Input layer: 2 units (corresponding to the first two principal components of the stimulus)
- Hidden layer 1: 64 units with ReLU activation
- Hidden layer 2: 64 units with ReLU activation
- Output layer: 1 unit (predicted mean spike count)

#### Training

The model was trained in PyTorch using standard supervised learning. Input features consisted of the first two principal components (PC_1_, PC_2_) of the stimulus representation, with empirical mean spike counts as target outputs. The principal components were standardized using *z*-score normalization (zero mean, unit variance) via scikit-learn’s `StandardScaler` to facilitate training convergence. Data were randomly partitioned into training (80%) and testing (20%) sets using a fixed random seed (seed = 42) for reproducibility. The model was trained for 200 epochs with the Adam optimizer (learning rate = 0.001), using mean squared error (MSE) between predicted and observed mean spike counts as the loss function. Training was performed in batch mode using the entire training set per epoch. Validation loss on the held-out test set was monitored every 20 epochs to assess convergence and detect potential overfitting.

#### Prediction

After training, the fitted model was evaluated on a regular 100 × 100 grid spanning the principal component space (PC_1_, PC_2_ ∈ [ − 10, 10]). Standardized coordinates for each grid point were passed through the trained network to generate predicted mean spike counts, producing a continuous representation of the tuning curve across the two-dimensional stimulus space. Gradients of the tuning curve with respect to the input features, ∇_**x**_*f* (**x**), were computed numerically and subsequently used for Fisher information calculations in the local information analysis.

The distribution of the first two principal components (PC1 and PC2) from the three-dimensional projection of natural images was well approximated by a two-dimensional Gaussian centered at zero. Accordingly, the stimulus prior *p*(**x**) was fitted as a zero-mean bivariate Gaussian with covariance matrix Σ.

#### Fisher Information

Fisher information was computed to quantify the encoding precision of neural populations. For each neuron, the tuning curve *f* (*x*) describes the mean firing rate as a function of the stimulus, and the gradient of the tuning curve *f* ^*′*^(*x*) was computed as the derivative with respect to the stimulus.

Under the assumption of conditionally independent Poisson spiking, the linear Fisher information of the population—which neglects the variance’s stimulus dependence—was calculated as:

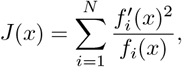

where *f*_*i*_(*x*) and 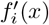 are the mean firing rate and its derivative for neuron *i*, respectively, and *N* is the number of neurons in the population. The contributions from all neurons are summed to obtain the population Fisher information for each stimulus.

#### Case of motion direction stimuli

In the case of the simulated direction-selective neurons, *f* (*x*) and *f* ^*′*^(*x*) were evaluated in practice at a grid of 50 evenly spaced stimulus values *x* in the interval [0, 2*π*]. The derivative *f* ^*′*^(*x*) was computed analytically. Instead, for the DS RGC quadruplets, for numerical stability, the tuning curve *f*_*i*_(*x*) of each neuron was first smoothly interpolated using a quadratic univariate spline (*k* = 2) with no smoothing (*s* = 0) to obtain a continuous representation across the stimulus space. The derivative 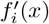 was then computed numerically from the spline representation at each of 180 stimulus positions uniformly distributed over the interval [0, 2*π*).

#### Case of natural image stimuli

For the model of RGC responses to natural images, the linear Fisher information is a matrix computed at each stimulus location as:

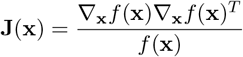

where *f* (**x**) is the predicted mean spike count and ∇*f* (**x**) is its gradient. ∇*f* (**x**) was estimated numerically using finite-difference approximation with step size *η* = 10^−4^, on a 100 × 100 grid spanning the images’ two-dimensional principal component space over the interval [−10, 10].

#### Precision

The mean squared error (MSE) of a Bayesian decoder, representing an ideal observer, was estimated at each stimulus location *x* using a Monte Carlo sampling procedure with 10,000 iterations (56; 11). At each iteration, neural responses **r** were generated by drawing Poisson spike counts with rate parameters given by the tuning curves evaluated at all stimulus positions x. The posterior probability of stimulus *x* given response **r**, *p*(*x*|**r**), was computed using Bayes’ rule:

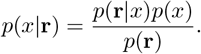

where the likelihood *p*(**r** | *x*) was calculated as the product of probabilities across a population of *N* neurons with independent Poisson spike counts **r** = (*r*_1_, *r*_2_, …, *r*_*N*_) given stimulus *x*. The likelihood is:

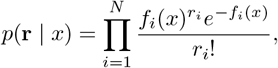

with *f*_*i*_(*x*) is the expected spike count of neuron *i* (its tuning curve) for stimulus *x*.

The marginal probability *p*(**r**) was obtained by integrating the joint distribution *p*(**r**, *x*) over the stimulus space using numerical integration (step size dx). To maintain numerical stability, probability distributions were log-transformed, and log-sum-exp calculations were used throughout.

Decoding performance was quantified by the circular mean squared error between the true stimulus and the decoded estimate. At each iteration, the decoded stimulus estimate 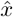 was obtained using the atan2 function, as the circular mean of *p*(*x* | **r**), the posterior probability of the stimulus given the sampled neural response **r**:

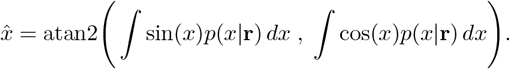

The circular squared error was calculated as:

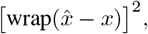

where `wrap()` denotes the wrapping operation to the interval [ − *π, π*]. The MSE was accumulated as a running average across all Monte Carlo samples and decoding precision as its inverse.

### Stimulus-specific information

#### Case of motion direction stimuli

In (Fig. 1 and 2), the stimulus-specific information (*I*_*SSI*_) and precision (the inverse of the mean squared decoding error) were estimated using a Monte Carlo sampling procedure with 10,000 iterations (56; 11). They were accumulated as a running average across samples, where the expectation is approximated by the sample mean.

The *I*_*SSI*_ was computed as:

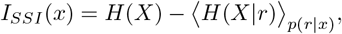

where *H*(*X*) represents the entropy of the stimulus distribution and *H*(*X* | *r*) is the conditional entropy of the stimulus ensemble *X* given the particular neural response *r*.

The stimulus entropy *H*(*X*) was calculated with a uniform prior probability of *N* = 180 equidistant stimuli, yielding

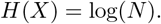

The conditional entropy *H*(*X*|**r**) was estimated as:

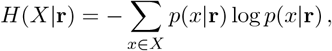

where the step size *dx* = 0.005 radians and the posterior probability *p*(*x* | **r**) was computed using Bayes’ rule, as for the calculation of the MSE:

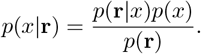

#### Case of natural image stimuli

For the RGC responses to natural image 2D projections in Fig. 3, stimulus-specific information (*I*_*SSI*_) was calculated via exhaustive enumeration of discretized stimulus and response spaces. In that case, *I*_*SSI*_ quantifies the information that a neural response provides about the stimulus presented at a specific location **x** in the two-dimensional principal component space.

The stimulus prior, *p*(**x**), was modeled as a zero-mean bivariate Gaussian with covariance matrix Σ_*x*_, fitted to the distribution of the first two principal components of natural images. Its entropy *H*(*X*) can be computed analytically as:

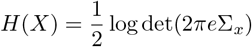

The conditional entropy *H*(*X*|*r*) was computed for each response *r* as:

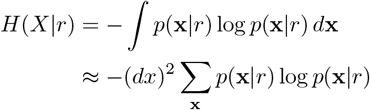

Computing *p*(**x** | *r*) involved brute-force evaluation over discretized stimulus and response spaces. The two-dimensional principal component space was discretized into a regular grid of 100 × 100 points with uniform spacing *dx* within the interval [−10, 10]. The space of possible spike counts was discretized as *r* ∈ {0, 1, …, *r*_max_}, where 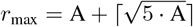 with A = 20, ensuring coverage of the tail of the Poisson distribution.

The log-probability log *p*(**x**) was evaluated at each grid point using the multivariate Gaussian formula, computing the Mahalanobis distance 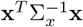 for each location. For each combination of stimulus location **x** and spike count *r*, the likelihood *p*(*r*|**x**) was computed using Poisson statistics:

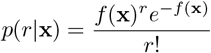

where *f* (**x**) is the predicted mean spike count from the neural network model. The posterior distribution *p*(**x**|*r*) was computed using Bayes’ rule:

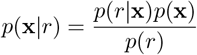

where the marginal likelihood *p*(*r*) was obtained by numerical integration over the stimulus space:

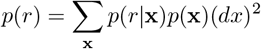

All computations were performed in log-space for numerical stability, using the log-sum-exp trick for the marginalization. The *I*_*SSI*_ was computed at each stimulus location **x** as:

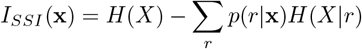

### Local information

#### Case of motion direction stimuli

In Figs. 1 and 2, local information was estimated via Monte Carlo sampling, using the following approximation for *I*_*local*_(*x*):

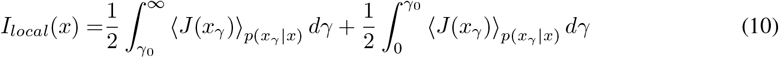

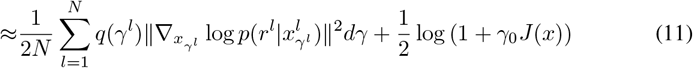

where the summation in the first term is taken over *N* samples, with *γ* is sampled from from an importance sampling distribution,

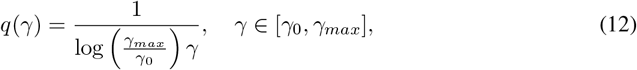

while *x*_*γ*_ is sampled from a wrapped gaussian:

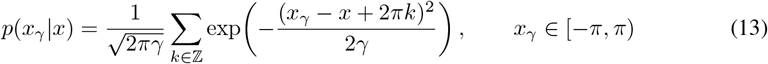

and *r* is sampled from *p*(*r*|*x*_*γ*_). We set *N* = 10^4^, *γ*_0_ = 0.01, and *γ*_max_ = 50.

#### Case of natural image stimuli

For a two-dimensional stimulus (Fig. 3), the decomposition of *I*_*local*_ takes the form:

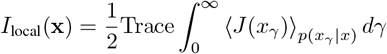

where *p*(*x*_*γ*_ | *x*) = 𝒩 (*x, γI*). To approximate this expression numerically, the integration over noise scales was partitioned into two regions: [0, *γ*_0_] and [*γ*_0_, *γ*_max_], where *γ*_0_ = 0.01 and *γ*_max_ = 200, as follows:

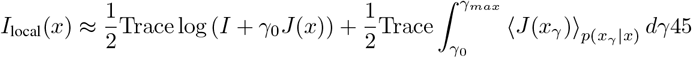

The integral over *γ* in the second term was approximated by computing the integrand at 50 increments of *γ*, which were equally spaced on log scale. Since we only considered 1 neuron, with discrete responses corrupted by Poisson noise, we could estimate the distribution, *p*(*r* | *x*_*γ*_), required to estimate the Fisher information *J* (*x*_*γ*_) numerically, by discretizing the 2-d stimulus space into equally spaced increments on a 2-d grid.

## 5 Acknowledgements

We thank F. Franke, M. Goldin and O. Marre for sharing the retinal data and S. Azeglio and T. Küen for insightful discussions. Ulisse Ferrari acknowledges this work was done within the framework of the PostGenAI@Paris project with the reference ANR-23-IACL-0007. Ulisse Ferrari benefited from financial support by the Agence Nationale de la Recherche (ANR) by grants NatNetNoise (ANR-21 CE37-0024), IHU FOReSIGHT (ANR-18-IAHU-01). Matthew Chalk and Steeve Laquitaine are funded by a grant RetNet4EC (ANR-22-CE92-0015), cofunded by the ANR and DFG. Our lab is part of the DIM C-BRAINS, funded by the Conseil Régional d’Ile-de-France. We thank Qube Research Technologies (QRT) for their financial support via the project DISSENSATION.

